# The Kinase CK1α coordinates Initiation and Termination of the cGAS-STING Pathway

**DOI:** 10.1101/2025.08.13.670063

**Authors:** Jane Jardine, Marine Tarrillon, Gwennan André-Grégoire, Kilian Trillet, Vanessa Josso, Laura Merlet, Rosalie Moreau, Luc Antigny, François Guillonneau, Alice Boissard, Cécile Henry, Joanna Re, Sophie Barillé-Nion, Nadine Laguette, Philippe P. Juin, Julie Gavard, Nicolas Bidère

## Abstract

The cGAS-STING pathway is an evolutionarily conserved antimicrobial defense mechanism that senses cytosolic DNA to trigger innate immune responses. cGAS and STING play dual roles in tumorigenesis, promoting antitumor immunity and cell death while fueling tumor growth and metastasis. However, the mechanisms fine-tuning this pathway remain elusive. Using proteomic approaches, we report that Casein Kinase 1 alpha (CK1α) operates as a bimodal regulator of the cGAS-STING pathway. CK1α supports optimal DNA sensing by preventing the proteasomal degradation of cGAS driven by the cullin-RING ubiquitin ligase 3 (CRL3). Conversely, CK1α facilitates STING degradation and signaling termination in response to STING agonists, tempering IRF3 activation. Exploiting these counterposing functions, we show that selective degradation of CK1α with molecular-glue degraders impaired the survival of a triple-negative breast cancer cell line with chronic cGAS-STING activation and synergized with a STING agonist to kill acute myeloid leukemia cells. Thus, CK1α’s dual regulatory role in the cGAS-STING pathway presents a promising target for therapeutic development.

**TEASER:** This study unveils CK1α as a bimodal regulator of the cGAS-STING pathway.

## INTRODUCTION

Aberrant cytosolic double-stranded DNA (dsDNA), resulting from pathogen infection or cellular stress, activates an evolutionarily conserved antimicrobial defense system orchestrated by the DNA sensor cyclic GMP-AMP synthase (cGAS) and its adapter protein Stimulator of Interferon Genes (STING). Upon binding to dsDNA, regardless of the sequence, cGAS undergoes liquid-liquid phase separation, forming liquid-like droplets that concentrate DNA and cGAS and favor the production of a cyclic dinucleotide 2′3′-cyclic GMP–AMP (cGAMP) (*1*, *2*). This secondary messenger then binds and activates the ER-resident adaptor protein STING, which shuttles to the Golgi to assemble a signaling platform with noncanonical IκB kinases (IKKs), TBK1 (tank-binding kinase 1), and IKKε (inhibitor of NF-κB kinase subunit ε) and transcription factors such as IRF3 (interferon regulatory factor 3) (*3–6*).

The phosphorylation of STING and IRF3 by TBK1/IKKε engages a broad transcriptional program that includes NF-κB-dependent proinflammatory cytokines and type I interferons (IFNs), leading to the expression of numerous interferon-stimulated genes (ISGs) and the efficient clearance of the infection (*1*). Of note, cGAS and STING have been linked to additional cellular processes, including metabolic homeostasis, life-and-death decisions, autophagy, and genome integrity (*1*). For instance, STING-mediated IRF3 signaling promotes cell-intrinsic apoptosis in an interferon alpha/beta receptor (IFNAR)-independent manner (*7*, *8*). Therefore, the tight regulation of cGAS and STING is crucial, and the underlying mechanisms are not fully understood.

Beyond its classical role in antimicrobial defense, the cGAS-STING pathway has emerged as a critical dual player in cancer immunology (*9*). For instance, chronic cGAS-STING signaling resulting from chromosomal instability (CIN), micronuclei, or DNA damage, whether endogenous or induced by chemotherapy (*10*), can foster an immunosuppressive, pre-metastatic environment (*9*). CIN, *i.e.* chromosome segregation errors, is a classical hallmark of cancer observed in over 60% of cancer cells and is associated with poor prognosis (*10*). CIN often leads to the formation of fragile micronuclei prone to breakage and leakage of DNA into the cytosol, acting as a genomic source of innate immune activation (*11–13*). Cancer cells displaying CIN, including triple-negative breast cancer (TNBC) cells, can develop an addiction to this chronic inflammation, and deleting cGAS and STING impairs their survival (*14*). In contrast, acute activation of the cGAS-STING pathway by anti-mitotic chemotherapy in breast cancer triggers paracrine pro-death signals (*15*). Moreover, controlled activation of the STING pathway can enhance anti-tumor immunity by directly eliminating malignant cells and promoting the recruitment and activation of adaptive immune cells. Accordingly, small-molecule agonists of STING have shown therapeutic efficacy against poor-prognosis hematologic human malignancies, including acute myeloid leukemia (AML). Given these dual roles, a key challenge lies in harnessing the cGAS-STING pathway to potentiate its anti-tumoral activity while mitigating its pro-tumorigenic effects.

CK1α is a pleiotropic kinase that regulates critical cellular pathways, including the developmental Wnt/β-catenin signaling pathway, lymphocyte activation, p53 regulation, programmed cell death, and autophagy (*16*, *17*). CK1α is crucial for the survival of aggressive hematologic human cancers and the immunomodulatory imide drug lenalidomide, which selectively connects CK1α to the cullin-RING ubiquitin ligase 4 complex (CRL4) to drive its ubiquitination and degradation, has proven efficacy in preclinical AML models (*18*, *19*). Nonetheless, the full landscape of CK1α functions remains ill-defined, in part due to the embryonic lethality of CK1α knockout models (*20*). Here, combining proteomic approaches, we report that CK1α functions as a “dual-gating” regulator of the cGAS-STING DNA-sensing pathway. First, CK1α gatekeeps the DNA sensor cGAS by maintaining its protein abundance, a process involving the CRL3 E3 ubiquitin complex. Second, CK1α accelerates the degradation of STING and thus tempers IRF3 signaling. We further show that targeting CK1α with molecular glue degraders counteracts chronic activation of cGAS-STING and cell survival in a chromosomally unstable TNBC cell line and potentiates the antitumor efficiency of STING agonists against AML models. Overall, our study highlights that CK1α is an important immunomodulator of the cGAS-STING pathway that could be of therapeutic interest.

## RESULTS

### CK1α Kinase is required for optimal cGAS Signaling

To define the proximity landscape interaction of CK1α, we employed an APEX-based proteomic strategy. To this end, we first engineered THP-1 monocytic cells deleted for endogenous expression of CK1α with single guide RNAs (CK1α^sg^), followed by lentiviral reconstitution with APEX2-tagged CK1α. Consistent with CK1α’s role in promoting the degradation of β-Catenin (*16*, *17*), CK1α^sg^ cells displayed elevated levels of β-Catenin, and this was effectively reversed upon expression of CK1α-APEX2 (**Fig. 1A and S1A**). Under these conditions, we identified 3,718 proteins as part of the CK1α proximity network (Log2 Ratio H_2_O_2_/Ctrl > 0,585; qValue < 0.05) from 4,209 detected proteins. As expected, the CK1α network was enriched with known substrates and binding partners, including FAM83 proteins – subcellular anchors of CK1α (*21–23*) – and key regulators of Wnt/β-catenin pathway (*16*, *17*) (**Fig. S1B**). We also detected proteins reported to dynamically associate with CK1α upon stimulation of antigen receptors (*24*, *25*), during autophagy (*26–29*), and in response to TNF receptor signaling (*30*, *31*), suggesting proximity in resting conditions (**Fig. S1B**). Notably, the non-canonical IKK Serine/threonine-protein kinase (TBK1), an essential kinase involved in regulating type I interferon (IFN) production during innate immune responses, was among the top-scoring hits (highest qValue) (**Fig. 1B**) (*5*, *32*). Gene Ontology (GO) analysis for positive regulators of type I IFN signaling (GO:0032481) further revealed enrichment of members of RIGI-MAVS, cGAS-STING, and TLR4-TRIF pathways within CK1α-proximity interactors (**Fig. 1C and 1D**). These three Pattern Recognition Receptors (PRR) and their adaptor proteins sense cytosolic DNA, viral RNA, and bacterial wall lipopolysaccharide (LPS), respectively, to elicit a proinflammatory immune response (**Fig. 1E**). To assess the functional importance of CK1α, CK1α^sg^ THP-1 cells were treated with selective agonists of cGAS (Y-shaped DNA G3-YSD), RIG-I (5’ triphosphate hairpin RNA 3p-hpRNA), or TLR4 (rough-form LPS-EK). Although the induction of interferon beta (IFNB1) mRNA was normal in response to RNA, we observed a diminished response to LPS and DNA, with a stronger inhibition observed with the latter (**Fig. 1F**). Similar results were obtained with two additional DNA species, *ie* an 80 bp synthetic dsDNA sequence and herring testes DNA (HT-DNA), a naturally derived DNA typically ranging from ∼400-1000 bp. Similar results were obtained with a second THP-1 sgRNA transduced clone (**Fig. 1G-I**). Moreover, CK1α deletion also abrogated the IFN-stimulated gene (ISG) CXCL10 expression and the NF-κB-induced IL6 expression (**Fig. 1J-K**). Hence, CK1α positively regulates type I IFN production upon DNA stimulation.

**Fig. 1.**
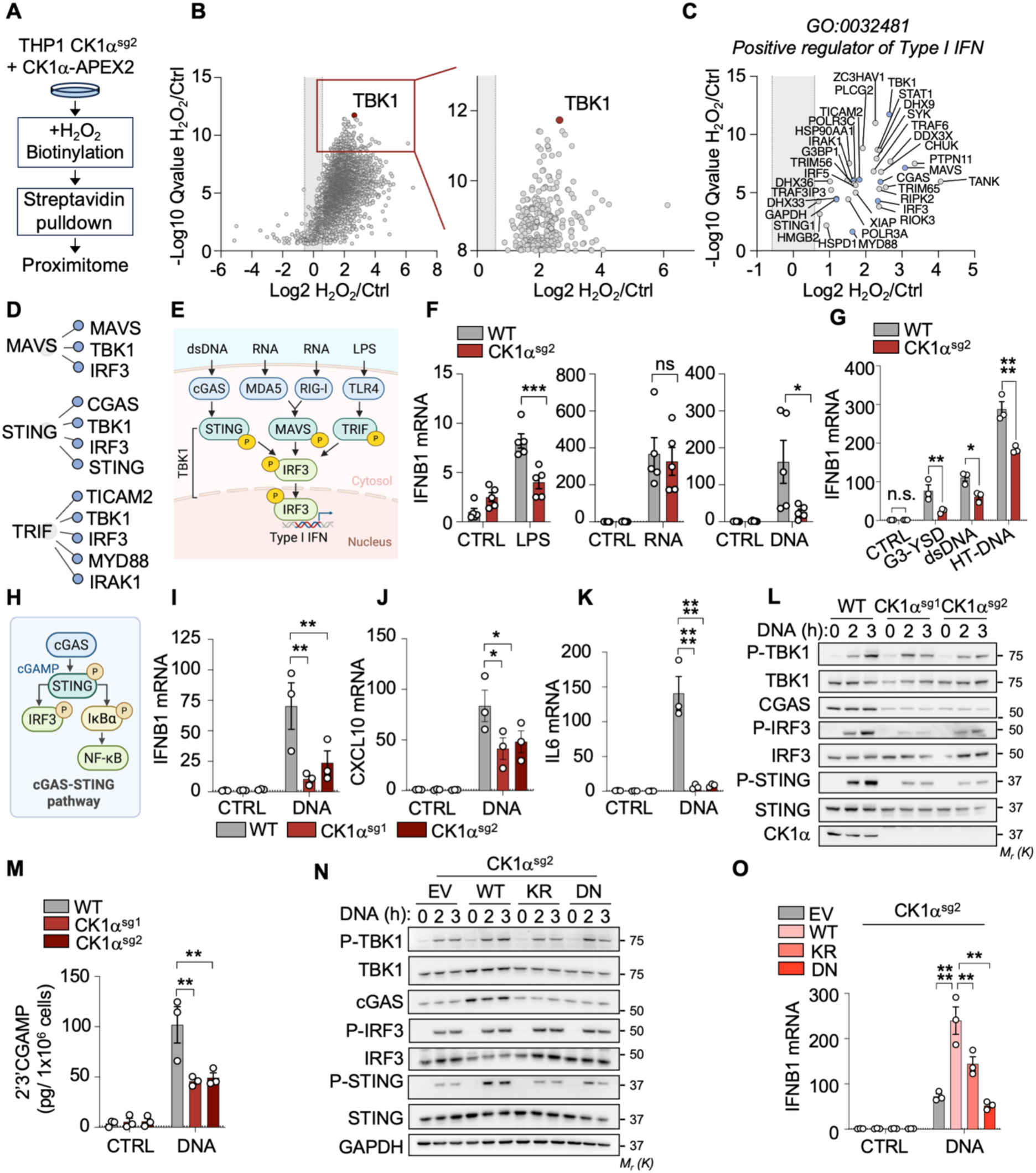
CK1α Kinase is required for optimal cGAS Signaling. (**A**) Schematic flow chart of the CK1α-APEX2-proximity-based proteomic assay. (**B**) Characterization of the global and steady state CK1α-proximity interaction landscape in sgRNA transduced THP-1 cells. (**C-D**) Gene Ontology analysis of positive regulators of Type I IFN (GO:0032481) within CK1α-proximity interactors. (**D**) Illustration of the members of the MAVS, STING, and TRIF pathways that are part of the CK1α proximity landscape (blue). (**E**) Schematic representation of the TBK1 substrates within the cGAS-STING, RIG-I/MDA5-MAVS, and TLR4-TRIF pathways. (**F-G**) IFNB1 expression in CK1α sgRNA transduced THP-1 cells measured by quantitative PCR with reverse transcription (RT-qPCR) after treatment with 1µg/mL G3-YSD (DNA) (F), 1 µg/mL 3p-hpRNA (RNA), and 1 µg/mL LPS-EK (LPS) for 2h or with 1 µg/mL G3-YSD, 1µg/mL of an 80 bp synthetic dsDNA sequence (dsDNA), and 1µg/mL of herring testes DNA (Ht-DNA) for 2h (G). (**H**) Simplified schematic of the signaling downstream of cGAS activation. (**I-K**) IFNB1, CXCL10, and IL6 expression quantified by RT-qPCR in THP-1 transduced with sgRNA targeting CK1α, after treatment with 1 µg/mL G3-YSD (DNA) for either 2h (I) or 6h (J-K). (**L and M**) Analysis of the TBK1-mediated phosphorylation cascade by immunoblotting (L) and measurement of cytosolic cGAMP by ELISA in sgRNA transduced THP-1 cells after treatment with 1µg/mL G3-YSD (DNA) for the indicated time (M). (**N-O**) THP-1 cells transduced with CK1α sgRNA were reconstituted with WT and catalytically dead CK1α (KR, DN) and treated for the indicated time with 1µg/mL G3-YSD (DNA). The phosphorylation of TBK1, IRF3, STING, and IκBα was assessed by immunoblotting (N), and the expression of IFNB1 was measured by RT-qPCR (O). Ordinary two-way ANOVA, *p < 0.05, ** p < 0.01, *** p < 0.001, **** p < 0.0001.

Downstream of DNA sensing by cGAS, TBK1 phosphorylates itself, STING, and IRF3 to drive type I IFN production (*5*). In CK1α^sg^ cells, the phosphorylation of STING and downstream activation of the cGAS-STING pathway were markedly reduced following transfection with synthetic DNA (**Fig. 1L**). This was also true in cells stimulated with the ABT-737 and S63845 BH3 mimetics, which cause mitochondrial outer membrane permeabilization and release of mitochondrial DNA into the cytosol (*33*, *34*) (**Fig. S1C**). We next measured the production of intracellular 2’3’-cGAMP by ELISA as a surrogate for cGAS activity (*1*, *2*), and found a decrease in its cyclic-GMP-AMP synthase activity in CK1α sgRNA transduced cells (**Fig. 1M**). Thus, CK1α regulates DNA sensing, either upstream or at the level of cGAS.

To determine whether CK1α’s catalytic activity is required, we inserted point mutations in its ATP pocket (K46R, D136N) to generate kinase-dead mutants (*35*, *36*). Reconstitution of CK1α-depleted cells with wild-type CK1α, but not CK1α K46R or D136N, restored DNA-induced signaling and IFNB1 induction (**Fig. 1N and 1O**). Thus, the kinase activity of CK1α is essential for promoting cGAS-STING pathway activation.

### CK1α controls the Turnover of cGAS

When comparing the CK1α proximity landscape at a steady state and after DNA stimulation, we did not observe overt change in the type I IFN signature (**Fig. 2A-B and S1D**). However, while examining the cGAS-STING signaling pathway by immunoblot, we noticed that cGAS protein levels were reduced in the absence of CK1α (**Fig. 1L, 1N, 2C**). Reintroduction of wild-type CK1α restored the abundance of cGAS, whereas a catalytically dead CK1α mutant (K46R) failed to do so (**Fig. 2E**). Notably, treatment with the molecular glue degraders DEG-77 and SJ3149, which are lenalidomide derivatives that selectively induce proteasomal degradation of CK1α (*18*, *37*), also led to reduced cGAS levels (**Fig. 2D**). In contrast, cGAS mRNA expression remained unchanged without CK1α, suggesting a post-translational regulation (**Fig. 2F**). Accordingly, chemical inhibition of the proteasome with bortezomib partially restored cGAS levels in CK1α sgRNA transduced cells (**Fig. 2G**). Thus, the CK1α kinase protects cGAS from proteasomal degradation.

**Fig. 2.**
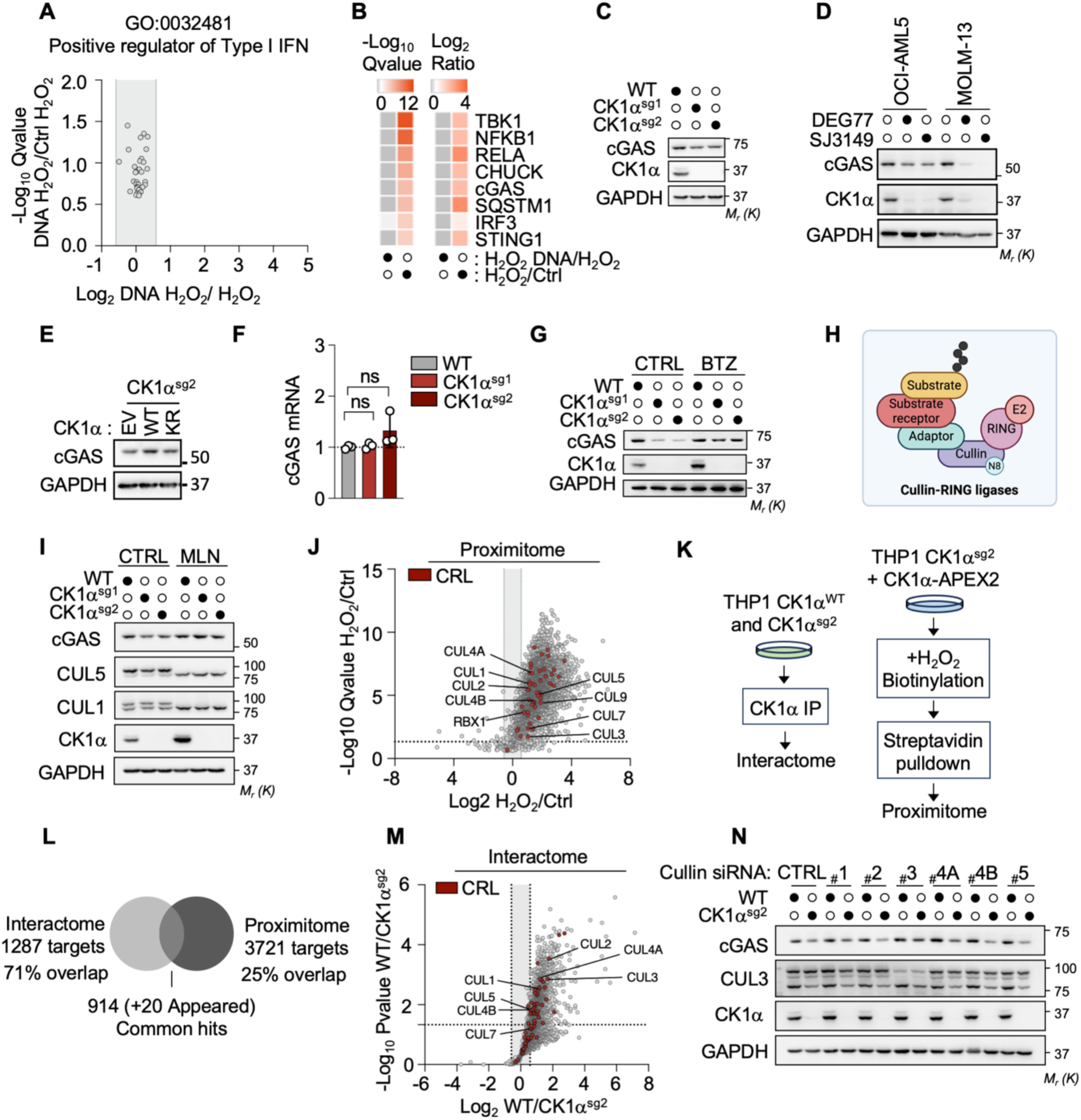
CK1α counteracts the Cullin-RING Ubiquitin Ligase-mediated Degradation of cGAS. (**A-B**) Characterization of the CK1α-APEX2 proximity landscape in THP-1 cells stimulated after 2 hours with 1µg/mL G3-YSD (DNA), normalized to the untreated condition. (**A**) Gene Ontology analysis of positive regulators of Type I IFN (GO:0032481) within the DNA-stimulated CK1α proximity landscape (H_2_O_2_ DNA/ H_2_O_2_). (**B**) Analysis of the core cGAS-STING pathway proteins (cGAS, STING, TBK1, IRF3, NF-κB, CHUCK, RELA, SQSTM1) within the CK1α proximity landscape. (**C**) cGAS protein abundance assessed by immunoblot in CK1α sgRNA transduced THP-1 cells. (**D**) cGAS protein abundance assessed by immunoblot in HL-60, OCI-AML5 and MOLM-13 pre-treated with 10 µM Q-VD-OPh for 15 Min and further treated with 100 nM DEG-77 or SJ3149 for 120h. (**E**) Immunoblotting analysis of CK1α sgRNA transduced THP-1 cells reconstituted with either an empty vector (EV), CK1α ^WT^ (WT), or catalytically dead CK1α^K46R^ (KR). (**F**) cGAS expression quantified in CK1α sgRNA transduced THP-1 cells by RT-qPCR. **(G)** cGAS protein levels assessed by immunoblot of CK1α sgRNA transduced THP-1 cells treated with the proteasome inhibitor Bortezomib (BTZ, 10 nM, 16h). (**H**) Illustrative representation of the core subunits of a typical CRL complex. **(I)** cGAS protein levels assessed by immunoblot of CK1α sgRNA transduced THP-1 cells treated with the protein neddylation inhibitor MLN492 (MLN, 24h, 1 µM). (**J**) Volcano plot highlighting the Cullin-RING Ligases (CRLs, in red) (GO: 003146) within the CK1α proximity landscape. (**K**) Schematic flow chart of the dual proteomic approach, CK1α-IP proteome (interactome) and CK1α-APEX2 proximity labeling proteome (proximitome) performed in THP-1. (**L**) Schematic visualization of the protein overlap between the CK1α-interactome and the proximitome performed in THP-1. (**M**) Volcano plot highlighting the Cullin-RING Ligases (CRLs, in red) (GO: 003146) within the CK1α interactome. (**N**) cGAS protein abundance assessed by immunoblotting in sgRNA transduced THP-1 cells transfected with the indicated Cullin targeting siRNA. Ordinary two-way ANOVA, *p < 0.05, ** p < 0.01, *** p < 0.001, **** p < 0.0001.

Cullin-RING ligases (CRLs), one of the largest superfamilies of ubiquitin E3 ligases (*38*), account for nearly 20% of protein degradation by the proteasome (*39*). CRLs are modular platforms built on a cullin scaffold (CUL1-CUL9) that partner with a dedicated RING finger protein (RBX1 or RBX2) and interchangeable substrate receptors to specifically recruit target proteins to be ubiquitinated (*40*) (**Fig. 2H**). The reversible binding of NEDD8 to cullins, a process called neddylation, is central to CRL activity (*39*). We found that treating cells with the neddylation inhibitor MLN492 stabilized cGAS protein levels in CK1α-depleted cells, suggesting a role for CRLs (**Fig. 2I**). Additionally, RBX1 and all eight cullin family members were found in the APEX2 proximity labeling proteome of CK1α (proximitome) (**Fig.1J**). To further analyze the interaction landscape of CK1α, we performed endogenous CK1α immunoprecipitation followed by proteomic analysis (**Fig. 2K-M**). This approach identified 1,287 CK1α-associated proteins, with 71% overlap between the CK1α-IP proteome (interactome) and CK1α-APEX2 proximity labeling proteome (proximitome) datasets, among which were the canonical cullins (CUL1-5) (**Fig. 2M**). Recently, CUL5 has been linked to the degradation of nuclear cGAS together with RBX2 and the substrate receptor SPSB3 (*41*). However, siRNA-mediated depletion of RBX2, CUL5, and SPSB3 did not prevent cGAS degradation in CK1α^sg^ cells, suggesting the involvement of an alternative CRL (**Fig. S2A**). Therefore, we next conducted a targeted siRNA mini-screen against canonical cullins (CUL1-CUL4B) and found that CUL3-targeted siRNA recovered cGAS protein levels (**Fig. 2N**). Thus, CK1α counteracts cGAS degradation through a CRL complex involving CUL3.

### CK1α Degraders enhance Paclitaxel-induced Cell Death in a TNBC Cell Line with chromosomal Instability

We next reasoned that targeting CK1α could mitigate the constitutive activation of the cGAS-STING pathway and increase the vulnerability of cancer cells displaying CIN. To test this hypothesis, we employed the TNBC BT-549 cell line, which is characterized by the presence of micronuclei and persistent cGAS-STING-dependent inflammatory signaling (*14*) (**Fig. 3A-B**). As expected (*14*), the use of a PROTAC degrader of TBK1 to suppress the cGAS-STING pathway significantly reduced cell viability (**Fig. S2B**). Treatment of BT-549 cells with three CK1α molecular glue degraders (BTX161, DEG-77, and SJ3149) for 48 hours led to a marked reduction of chronic inflammation signaling, including reduced constitutive phosphorylation of IRF3 and decreased expression of IRF3-and NF-κB-dependent cytokines (**Fig. 3C-F**). This was accompanied by a significant reduction in cell viability (**Fig. 3G**). Moreover, both DEG-77 and SJ3149 enhanced the cell-killing effect of Paclitaxel, a standard-of-care taxane for TNBC (*42*), and scored positively in the highest single-agent (HSA) synergy score (**Fig. 3G-H**). Thus, targeting CK1α with molecular glue degraders interferes with chronic inflammation associated with CIN and synergizes with Paclitaxel in a model of TNBC cells.

**Fig. 3.**
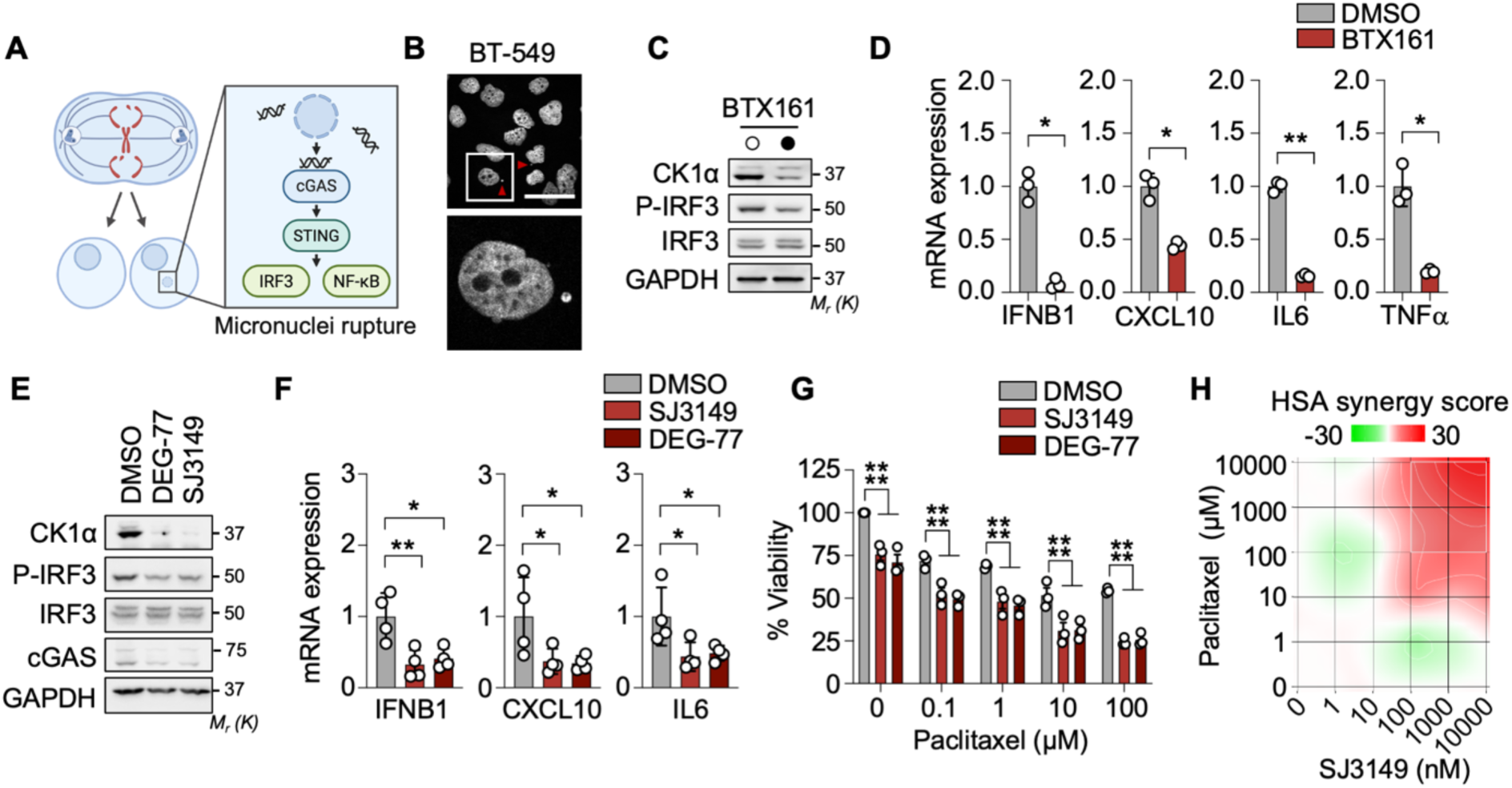
CK1α Degraders enhance Paclitaxel-mediated Cell Death in a TNBC Cell Line with chromosomal Instability. (**A**) Schematic illustration of micronuclei formation upon CIN and downstream cGAS-STING signaling. (**B**) BT-549 cells were stained with DAPI and visualized by confocal microscopy immunofluorescence. The micronuclei are indicated with a red arrow. Scale bar: 50 µm. (**C-D**) BT-549 cells were treated for 48h with 10 µM of the CK1α-targeting molecular glue BTX161. Activation of the cGAS-STING pathway was analyzed by immunoblotting (P-IRF3) (C) and RT-qPCR (IFNB1, CXCL10, IL6, and TNFα) (D). (**E-F**) BT-549 cells were treated with 1 µM of the CK1α-targeting molecular glues DEG-77 or SJ3149 for 48h. The activation of the cGAS-STING pathway was analyzed as in D-E. (**G**) Cell viability of BT-549 cells pre-treated with SJ3149 or DEG-77 (1 µM, 48h) and subsequently co-treated with escalating concentrations of Paclitaxel (24h) was assessed by Celltiter Glo. (**H**) Synergy distribution maps calculated with the highest single agent (HSA) model from BT-549 cells pre-treated with SJ3149 (24h, indicated concentrations) and further co-treated with Paclitaxel (24h, indicated concentrations). Ordinary two-way ANOVA, *p < 0.05, ** p < 0.01, *** p < 0.001, **** p < 0.0001.

### CK1α tempers STING Signaling

To explore if CK1α also plays a role in the cGAS-STING downstream of cGAS, we employed STING agonists (diABZI and cGAMP) to bypass cGAS (*43*). Unexpectedly, CK1α-depleted cells displayed increased phosphorylation of TBK1, IRF3, and STING, combined with enhanced expression of IFNB1 and downstream ISGs (**Fig. 4A-D**). Likewise, the siRNA-based silencing of CK1α or its degradation with the molecular glue degrader BTX161 boosted the phosphorylation cascade downstream of STING and increased IFNB1 mRNA expression (**Fig. S3A-C**). The enhanced STING signaling was associated with delayed STING degradation in CK1α^sg^ cells upon diABZI stimulation (**Fig. 4C**). Thus, CK1α has a dual-gating function on the cGAS-STING pathway, with a positive role in the maintenance of cGAS abundance and a negative role in signaling downstream of STING.

**Fig. 4.**
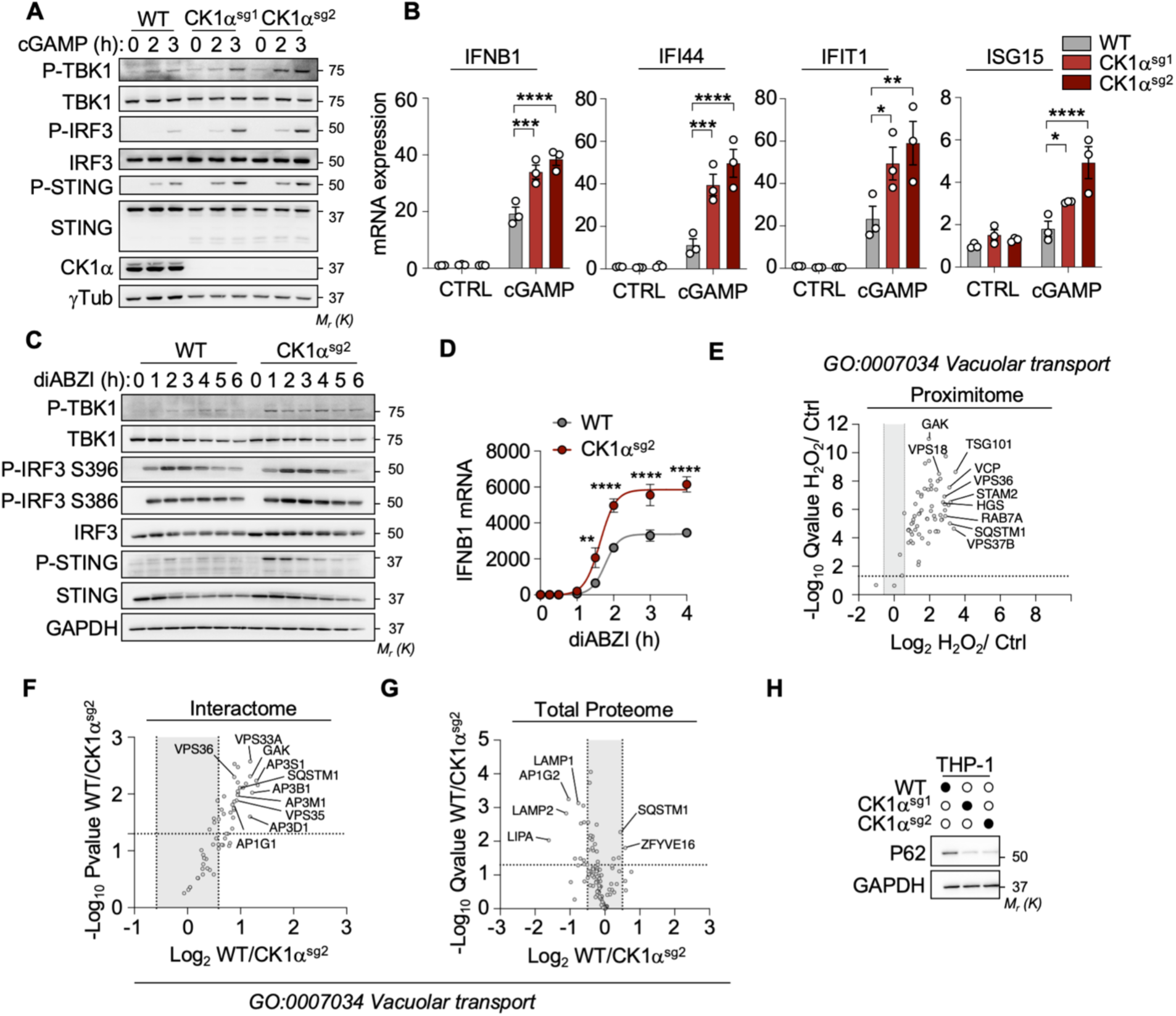
CK1α tempers STING Signaling. (**A**) Immunoblots of cell lysates from THP-1 cells transduced with sgRNA against CK1α and stimulated with the STING agonist 3’3’-cGAMP (10 µg/ml) for the indicated time. (**B**) IFNB1 and ISG expression levels quantified by RT-qPCR in sgRNA transduced THP-1 cells treated for 2h (IFNB1) or 4h (ISGs) as in (A). (**C-D**) Immunoblots of cell lysates (C) and IFNB1 expression quantified by RT-qPCR (D) in sgRNA transduced THP-1 stimulated with the STING agonist diABZI (2.5 µM) for the indicated time. (**E-G**) Gene Ontology analysis of Vacuolar Transport proteins (GO:0007034) within the CK1α proximitome (E), interactome (F), and total proteome (G). (**H**) Immunoblots of cell lysates from CK1α-sgRNA transduced THP-1 cells to asses p62 protein levels. Ordinary two-way ANOVA, *p < 0.05, ** p < 0.01, *** p < 0.001, **** p < 0.0001.

Following activation by cGAMP, STING traffics to the lysosome for degradation, thereby terminating signaling (*44–48*). This lysosomal targeting is mediated by clathrin-coated vesicles, the Endosomal Sorting Complex Required for Transport (ESCRT) complex, or autophagy mediated by the adapter protein p62 (SQSTM1) (*44–47*). Our proteomic datasets, focused on vacuolar transport proteins (GO:0007034), identified p62 in both the interactome and proximitome of CK1α, supporting previous findings (*26*) (**Fig. 4E and 4F**). Moreover, an unsupervised proteomic analysis of control and CK1α^sg^ cells using label-free quantitative proteomics unveiled a downregulation of p62 in the absence of CK1α (**Fig. 4G**), which was confirmed by immunoblotting analysis (**Fig. 4H**). Hence, CK1α regulates the abundance of the autophagy adaptor p62 and participates in optimal STING degradation upon stimulation.

### CK1α Degraders synergize with a STING Agonist to enhance Apoptosis in AML Cells

STING agonists have been shown to induce apoptosis in T cells and hematologic malignancies, including AML, by upregulating the proapoptotic BH3-only NOXA, PUMA, and BIM via TBK1 and IRF3, independently of TP53 (*7*, *8*). Accordingly, treatment with diABZI induced cell death across a panel of AML cell lines regardless of their TP53 status (**Fig. 5A**). In contrast, CK1α-targeted molecular glues (DEG-77 and SJ3149) efficiently degraded CK1α and reduced the viability of the TP53 wildtype AML cell lines only, supporting previous reports (*18*, *37*) (**Fig. 5A-B**). Given the regulatory role of CK1α in both STING and p53 signaling, we inferred that CK1α degraders could be combined with STING agonists to enhance apoptosis (**Fig.5C**). To test this, representative AML cell lines were treated with suboptimal concentrations of diABZI and the CK1α degraders DEG-77 or SJ3149. This drug combination killed AML cells more efficiently than monotherapy (**Fig.5D**). In line with this, DEG-77 and diABZI synergized to enhance the type I IFN and NF-κB response and the induction of the BH3-only *NOXA (PMAIP1)*, *PUMA (BBC3)*, and *BIM (BCL2L11)* (**Fig. 5E-G**). A similar trend was observed in TP53-deleted cell lines THP-1 and HL-60 (**Fig. S3D**). Collectively, these results highlight the potential therapeutic interest of CK1α-targeted molecular glues in combination with STING agonists.

**Fig. 5.**
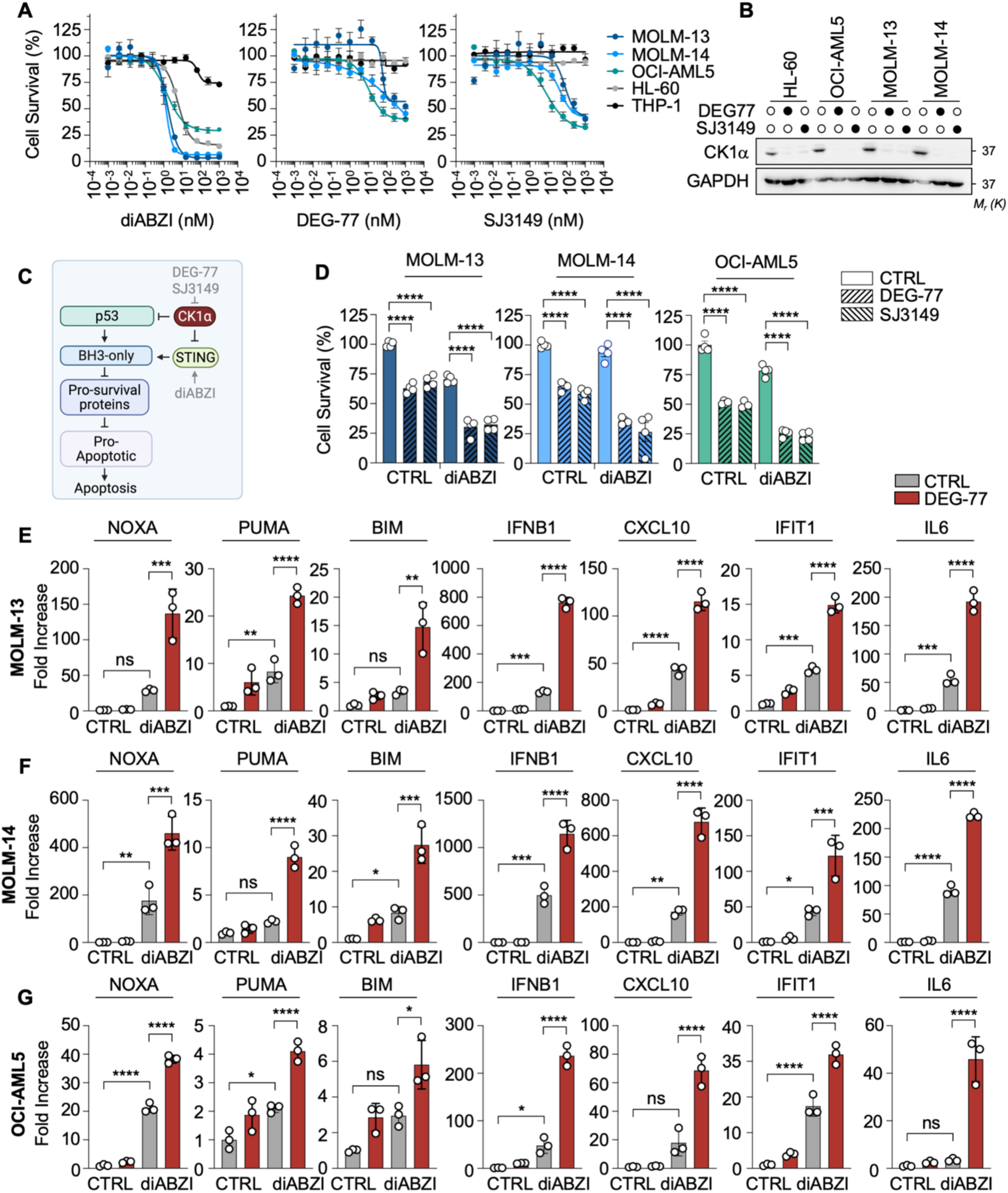
CK1α Degraders synergize with a STING Agonist to enhance Apoptosis of AML Cells. (**A**) Cell survival, assessed by Celltiter glo, in a panel of AML cells (TP53 mutated: THP-1, HL-60, and TP53 wildtype: OCI-AML5, MOLM-13, MOLM-14) upon diABZI, DEG-77, or SJ3149 titration over 24h. (**B**) The abundance of CK1α was assessed by immunoblot across a panel of AML cells upon treatment with the CK1α molecular glues DEG-77 and SJ3149 (1 µM, 24h). (**C**) Schematic representation of the apoptotic pathway induced downstream of CK1α inhibition. (**D**) Cell viability was assessed across a panel of p53 WT AML cells (MOLM-13, MOLM-14, OCI-AML5) upon pre-treatment with CK1α degraders (100 nM DEG-77 or 50 nM SJ3149) for 4h and stimulation with diABZI at 1.25 nM (MOLM-14), 2.5 nM (OCI-AML5, MOLM-13) for 16h. (**E-G**) Quantification of pro-apoptotic genes (NOXA, PUMA, BIM) and cGAS-STING target genes (IFNB1, CXL10, IFIT1, IL6) expression by RT-QPCR in TP53 wildtype AML cells: MOLM-13 (E), MOLM-14 (F), and OCI-AML5 (G) cells. To prevent apoptosis, cells were pre-treated with the pancaspase inhibitor Q-VD-OPh (10 µM, 15 Min), then treated with DEG-77 (100 nM, 4h) and further stimulated with diABZI (5 nM, 16h) prior to qPCR analysis. Ordinary two-way ANOVA, *p < 0.05, ** p < 0.01, *** p < 0.001, **** p < 0.0001.

## DISCUSSION

cGAS and STING play central roles in innate immune surveillance by detecting cytosolic double-stranded DNA from pathogens or damaged cells, initiating pro-inflammatory responses. This signaling pathway has also emerged as a critical regulator of cancer immunology, exhibiting a paradoxical influence on tumorigenesis (*9*). Chronic activation of this pathway resulting from chromosomal instability fosters tumor cell survival, while acute activation enhances anti-tumor immunity by eliminating malignant cells and mobilizing adaptive responses (*9*). This duality underscores the need for a deeper understanding of the molecular mechanisms governing the cGAS-STING pathway.

Our results have identified the kinase CK1α as a bimodal regulator of the cGAS-STING pathway, ensuring a tightly controlled balance between activation and restraint. This dual gating function is reminiscent of CK1α’s role in antigen receptor signaling, where it first allows signal transduction to NF-κB transcription factors and subsequently terminates signaling (*24*, *25*). Our combined analysis of the interactome and proximitome of CK1α captured several established partners and identified novel interactors that are likely to participate in regulating the cGAS-STING pathway. However, our experiments did not reveal inducible CK1α interactions with known components of the pathway upon DNA stimulation. Thus, whether CK1α is actively recruited to signaling complexes or exerts its gatekeeper function through constitutive regulation of proximal interactors remains to be determined. Here, we provide evidence that CK1α governs the abundance of cGAS by preventing its proteasomal degradation by a CRL3 complex. Nonetheless, future work is required to identify among the hundreds of substrate receptors (*40*), the one involved here. This expands the growing list of post-translational modifications – including phosphorylation (*49*), SUMOylation (*50*, *51*), ISGylation (*52*), acetylation (*53*, *54*), methylation (*55*, *56*), glutamylation (*57*), and ubiquitylation (*41*, *58–62*) – regulating key aspects of cGAS function, such as DNA binding, oligomerization, enzymatic activity, and protein stability. Moreover, our work echoes the recent description by Ablasser and colleagues of the CUL5-RBX2-SPSB3 CRL complex in the regulation of nuclear cGAS turnover to prevent aberrant activation of chromatin-bound cGAS following mitotic exit (*41*). Combined, these results suggest that different CRLs may be at play to tightly regulate cGAS depending on the cellular context.

In addition to protecting cGAS from proteasomal degradation, we found that CK1α marshals STING response to prevent excessive signaling. Our data show that CK1α stabilizes p62, a key adaptor protein involved in the autophagic clearance of STING (*45*). Accordingly, the overexpression or pharmacological activation of CK1α enhances p62-mediated STING degradation in mouse embryonic fibroblasts and mitigates type I IFN-driven inflammation (*26*). Moreover, the treatment with a CK1α activator drives CK1α to directly phosphorylate p62 to enhance STING targeting to autophagosomes (*26*), suggesting that CK1α not only governs p62 abundance but may also modulate its functional state. In addition to conveying inflammation, STING functions as a proton channel at post-Golgi vesicles (*63*, *64*), regulates non-canonical autophagy (*65–67*), and participates in lysosome biogenesis through TFEB/TFE3 expression (*67–69*). Our future work will aim to define whether CK1α’s regulatory influence extends to these newly defined branches of STING function.

From a therapeutic perspective, our study underlines the potential of exploiting the dual gating function of CK1α to modulate the cGAS-STING pathway in a context-dependent manner. First, we show that CK1α-directed molecular glues can be employed in a TNBC cell line displaying CIN to suppress aberrant chronic cGAS-STING activation. These compounds may also hold therapeutic potential in autoinflammatory syndromes marked by perturbed self-DNA metabolism involving cGAS and STING, such as in Aicardi-Goutières syndrome or DNase II deficiency, where type I IFN are elevated (*70*). Conversely, our study reports that these same molecular glues can be combined with STING agonists to potentiate proapoptotic STING signaling and enhance tumor cell killing. Acute STING signaling has indeed been reported to promote apoptosis of T cells and hematologic malignancies (*7*). Our proposed synergistic strategy aligns with the growing trend of combination therapy to maximize STING agonist efficacy. Such strategies include co-treatment with proapoptotic BH3 mimetics (*8*), CAR-T cells (*71–73*), immunotherapy with immune checkpoint inhibitors, or bispecific antibodies (*74–76*) and radiotherapy (*76*). Together, our findings establish CK1α as a crucial rheostat in cGAS-STING signaling, with implications for both immune regulation and cancer therapy.

## MATERIALS AND METHODS

### Cell culture and reagents

HEK-293T, HL-60, MOLM-14, OCI-AML5, and THP-1 cells were purchased from the American Type Culture Collection. MOLM-13 and the BT-549 cells were generously gifted by Christina Woo (Harvard University, US) and Floris Foijer (University of Groningen, Netherlands), respectively. Cells were cultured in Dulbecco’s modified Eagle’s medium DMEM (HEK-293T) or RPMI 1640 (THP-1, HL-60, MOLM-13, MOLM-14, OCI-AML5, BT-549) supplemented with 10% (HEK-293T, BT-549) or 20% (AML cell lines) FBS, 1% Glutamax, 100 mg/ml penicillin, 100 mg/ml streptomycin, and 50 µg/ml normocin (Invivogen). The culture media from AML cells was further supplemented with 1% non-essential amino acids, 1µM Hepes, and 100µM sodium pyruvate. The cells were maintained at 37°C in a humidified incubator with 5% CO_2_. All cells were regularly tested and found to be negative for mycoplasma contamination. Cells were treated as indicated with the following compounds: 3’3’-cGAMP Fluorinated (tlrl-nacgaf-05, Invivogen), 3p-hpRNA (tlrl-hprna, Invivogen), ABT-199 (S8048, Selleckchem), Bafilomycin (201550A, Santa Cruz), Bortezomib (S1013, Selleckchem), BTX161 (HY-120084, MedChemExpress), DEG-77 (HY-157334, MedChemExpress), diABZI (S8796, Selleckchem), G3-YSD (tlrl-ydna, Invivogen), HT-DNA (D6898, Sigma), LPS-EK (tlrl-peklps, Invivogen), MLN4924 (SI9830, LifeSensors), Paclitaxel (S1150, Selleckchem), Q-VD-OPh (S7311, Selleckchem), S638445 (S8383, Selleckchem) and SJ3149 (HY-160444, MedChemExpress). The TK1 PROTAC 3i (7259) and its control (7260) were from Tocris.

### Site-directed mutagenesis

The pLC-FLAG-CSNK1A1-K46R-Puro and pLC-FLAG-CSNK1A1-D136N-Puro plasmids were generated by PCR-mediated site-directed mutagenesis of the pLC-FLAG-CSNK1A1-WT-Puro plasmid (#123319, Addgene) using the following primers: CK1A-D136N forward, GAAGTTATCTGGTTTAAT<u>GTT</u>TCTGTGTATAAAATTC; CK1A-D136N reverse, GAATTTTATACACAGA<u>AAC</u>ATTAAACCAGATAACTTC; CK1A K46R forward, CTTCTGAGATTCTAG<u>CCT</u>CACTGCCACTTCCTC; CK1A K46R reverse, GAGGAAGTGGCAGTG<u>AGG</u>CTAGAATCTCAGAAG. The point mutations were verified by sequencing. pLC-Flag-CSNK1A1 WT-Puro was a gift from Eva Gottwein (Addgene plasmid # 123319; http://n2t.net/addgene:123319; RRID:Addgene_123319)

### Transduction of CK1α sgRNA and lentiviral transduction

THP-1 cells were transfected by electroporation with the pSpCas9(BB)-2A-GFP (PX458) - sgCSNK1A1 (sgRNA: 5’-TGTACTTATGTTAGCTGACC-3’) plasmid using the ECM 830 electroporation system (BTX Apparatus) (*25*). Briefly, 10×10^6^ THP-1 cells were electroporated at 260V for 10 ms with 10 µg of PX458-sgCSNK1A1 plasmid in a 4 mm electroporation cuvette (VWR). GFP-positive cells were single-cell sorted using a FACS Aria III (Cytocell core facility, SFR Bonamy, BioCore, Inserm UMS 016, CNRS UAR 3556, Nantes, France). The stable expression of the rescue constructs pLC-FLAG-CSNK1A1-WT-Puro, pLC-FLAG-CSNK1A1-K46R-Puro, and pLC-FLAG-CSNK1A1-D136N-Puro was performed by lentiviral infection in CK1α sgRNA transduced THP-1 cells as previously described. Briefly, lentiviral particles were produced in HEK-293T cells by transfecting them with the lentiviral packaging plasmids psPAX2 and pVSV-G. The particles were centrifuged at 1,000*g* for 90 min onto the THP-1 CSNK1A1 KO cells with 8 μg/ml polybrene (Santa Cruz). Infected cells were then selected using 1.25 µg.ml^-1^ puromycin.

### Stimulation with PRR agonists

THP-1 were transfected with 1 µg/ml G3-YSD (tlrl-ydna, Invivogen), 1 µg/ml 80nt dsDNA probe (Nadine Laguette, IGMM) (*77*), 1 µg/ml HT-DNA (D6898, Sigma) or 100 ng/ml 3p-hpRNA (tlrl-hprna, Invivogen) using Lipofectamine 2000 (Invitrogen) following the manufacturer’s recommendations.

DNA probe sequence: Forward, ACATCTAGTACATGTCTAGTCAGTATCTAGTGATTATCTAGACATACATGATCTATGACATATATAGTGGATAAGTGTGG ; Reverse, CCACACTTATCCACTATATATGTCATAGATCATGTATGTCTAGATAATCACTAGATACTGACTAGACATGTACTAGATGT The other agonists were directly added to the media: 10 µg/ml 3’3’-cGAMP (tlrl-nacgaf-05, Invivogen), 1 µg/mL LPS-EK (tlrl-peklps, Invivogen) or diABZI (S8796, Selleckchem) at the indicated concentrations.

### siRNA transfection

Silencing by siRNA in THP-1 cells was performed by electroporation. Briefly, 4×10^6^ THP-1 cells were washed in PBS1X, resuspended in 400 µL opti-MEM with 80pmol siRNA in a 4 mm electroporation cuvette (VWR), then were electroporated at 260V for 10 ms using the ECM 830 electroporation system (BTX Apparatus).

The following siRNA duplexes were used: Stealth nonsilencing control (low-GC 12935111; Life Technologies); CK1α, 5’-GGCAAGGGCUAAAGGCUGCAACAAA-3’; CUL1, 5’-GGUCGCUUCAUA AACAAC A-3’; CUL2, 5’-GAGCUAGCAUUGGAUAUGUGG-3’; CUL3 5’-GAAGGAAUGUUUAGGGAUA-3’; CUL4A, 5’-GAACUUCCGAGACAGACCU-3’; CUL4B, 5’-AAGCCUAAAUUACCAGAA A-3’; RBX1#A, SASI_Hs01_00066597 (Sigma); siRBX1 (si#B), SASI_Hs01_00066598 (Sigma); RBX2 (si#D), SASI_Hs02_00345291 (Sigma); CUL5, (L-019553-00-0005) ON-TARGET SMARTpool (Dharmacon); SPSB3, (L-017713-00-0005) ON-TARGET SMARTpool (Dharmacon).

### Micronuclei Immunofluorescence

BT-549 cells were seeded onto 12 mm diameter coverslips, fixed with 4% paraformaldehyde, then permeabilized with 0.2% Triton-×100. The coverslips were incubated with 3% Bovine serum albumin, stained with DAPI (1µg/ml), and mounted onto glass slides using the prolong gold antifade mounting reagent (Invitrogen).

Single plane images were acquired by confocal microscopy (Nikon A1rHD LFOV) with a 60x oil-immersion lens at the MicroPICell core facility (SFR Bonamy, BioCore, Inserm UMS 016, CNRS UAR 3556, Nantes, France). Images were analyzed with the ImageJ software.

### Quantification of cGAMP by ELISA

Cells were lysed in TNT buffer (50 mM Tris HCl pH 7.4, 150 mM NaCl, 2 mM EDTA, 1% Triton-×100, 1% NP-40) supplemented with 1X Halt Protease Inhibitor cocktail (ThermoFisher Scientific). Samples were cleared by centrifugation at 10,000*g,* and intracellular cGAMP was quantified from cell lysates using the 2’3’-cGAMP ELISA kit (Cayman Chemicals), following the manufacturer’s instructions.

### Immunoblotting and immunoprecipitation

Cells were lysed in RIPA buffer (25 mM Tris HCl pH 7.4, 150 mM NaCl, 0.1% SDS, 0.5% Na-Deoxycholate, 1% NP-40, 1 mM EDTA) supplemented with 1X Halt Protease Inhibitor cocktail (ThermoFisher Scientific). Insoluble material was removed by centrifugation at 10,000*g*. For immunoblotting, 10 µg of lysates were diluted into 2X Laemmli buffer (Life Technologies) containing 10% 2-β-Mercaptoethanol and denatured at 95°C for 3 min. For immunoprecipitation experiments, 1 mg of lysate was precleared with Protein G Sepharose (Sigma) for 30 min, before incubation with 1 µg of antibody and Protein G Sepharose for 2h at RT or overnight at 4°C under rotation. Samples were washed in lysis buffer four times before the addition of 2X Laemmli buffer and immunoblotting. The samples were resolved onto 3-15% Tris-Acetate gels using the NuPAGE system (Invitrogen) and transferred onto nitrocellulose membranes (GE Healthcare) for immunostaining. The following antibodies were obtained from Cell Signaling Technology: HA-Tag (3724), IκBα (9242), phospho-IκBα Ser32/36 (9246), IRF3 (4302), phospho-IRF3 Ser396 (4947), p62 (88588), STING (13647), phospho-STING ser366 (19781), TBK1 (3504), phospho-TBK1 Ser172 (5483). CK1α (ab223144, for western blotting), phospho-IRF3 Ser386 (ab76493), RBX2 (ab181986) were purchased from Abcam. CK1α (A301-991A, from immunoprecipitation), CUL5 (A302-173A), RBX1 (A303-462A) were ordered at Bethyl. CUL1 (SC-17775), GAPDH (SC-32233), γTubulin (SC-51715), and PARP (SC-8007) were distributed by Santa Cruz, β-catenin (610153) by BD Bioscience, and CUL3 (11107-1-AP) by Proteintech.

### RT-qPCR analysis

Prior to RNA extraction, BT-549 (2-5×10^6^) were treated for 48h with the CK1α molecular glue degraders (10 µM BTX161, 1 µM DEG-77, or 1µM SJ3149). The AML cells (1.5-3 ×10^6^ cells) were treated as indicated. RNA extractions were performed using the Macherey-Nagel RNA Plus NucleoSpin kit on biological triplicates. The RNA was subsequently reverse transcribed using the Maxima First Strand cDNA synthesis kit (Thermo scientific) and 25ng of the resulting cDNA was amplified by qPCR using PerfeCTa SYBR Green SuperMix Low ROX (QuantaBio). Data were analyzed using the 2-ΔΔCt method and normalized by the housekeeping genes ACTB and HPRT1. The following primers were used: ACTB forward, 5ʹ-GGACTTCGAGCAAGAG ATGG-3ʹ; ACTB reverse, 5ʹ-AGCACTGTGTTGGCGTACAG-3ʹ; HPRT1 forward, 5ʹ-TGACACTGGCAAAACAATGCA-3ʹ; HPRT1 reverse, 5ʹ-GGTCCTTTTCACCAGCAAGCT-3ʹ; IFNB1 forward, 5’-GCTTGGATTCCTACAAAGAAGCA-3’; IFNB1 reverse, 5’-ATAGATGGTCAATGCGGCGTC-3’ CXCL10 forward, 5’-GAAAGCAGTTAGCAAGGAAAGGTG-3’; CXCL10 reverse, 5’-ATGTAGGGAAGTGATGGGAGAGG-3’; IFIT1 forward, 5’-GCGCTGGGTATGCGATCTC-3’; IFIT1 reverse, 5’-CAGCCTGCCTTAGGGGAAG-3’; IFI44 forward, 5’-ATGGCAGTGACAACTCGTTTG-3’; IFI44 reverse, 5’-TCCTGGTAACTCTCTTCTGCATA-3’; ISG15 forward, 5’-CGCAGATCACCCAGAAGATCG-3’; ISG15 reverse, 5’-TTCGTCGCATTTGTCCACCA-3’; IL6 forward, 5’-GACCCAACCACAAATGCCAG-3’; IL6 reverse, 5’-GTGCCCATGCTACATTTGCC-3’; TNF forward, 5’-CCCAGGCAGTCAGATCATCTTCT-3’; TNF reverse, 5’-ATGAGGTACAGGCCCTCTGAT-3’; NOXA forward, 5’-ACCAAGCCGGATTTGCGATT-3’; NOXA reverse, 5’-ACTTGCACTTGTTCCTCGTGG-3’; PUMA forward, 5’-GACCTCAACGCACAGTACGAG-3’; PUMA reverse, 5’-AGGAGTCCCATGATGAGATTGT-3’; BIM forward, 5’-TAAGTTCTGAGTGTGACCGAGA-3’; BIM reverse, 5’-GCTCTGTCTGTAGGGAGGTAGG-3’

### Cell viability assay and synergy score

Cell viability and drug synergy were assessed using the CellTiter-Glo Luminescent assay (Promega) following the manufacturer’s recommendation. Experiments were performed in 96-well plates using 10,000 BT-549 cells in a final volume of 100 µL. For the CK1α degraders experiments, the cells were first pre-treated with 1 µM (or indicated concentration) of CK1α degraders (SJ3149, DEG-77) for 48h then co-treated with Paclitaxel (Selleckchem) at the indicated concentrations for another 24h. For the TBK1 PROTAC experiments, the cells were treated with 1 µM TBK1 or control PROTAC for 48h. The synergy score was calculated using the Synergy Finder (version 3.0)(*78*) with the Highest single agent (HSA) model.

Likewise, for AML cells (THP-1, HL-60, OCI-AML5, MOLM-13 and MOLM-14), experiments were performed in 96-well plates in a final volume of 100 µL. The diABZI, DEG-77, and SJ3149 titrations were performed over 24h.

Drug combination experiments were performed as follows. AML cells were pre-treated for 4h with 100 nM DEG-77 or 50 nM SJ3149 then co-treated with diABZI at 1.25 nM (MOLM-14), 2.5 nM (OCI-AML5, MOLM-13), 5 nM (HL-60) or 200 nM (THP-1) for 16h.

### Proteomics – whole proteome analysis and CK1α immunoprecipitation analysis

Control and CK1α sgRNA transduced THP1 cells (3×10^6^ cells) were centrifugated for 3 min at 500g. The pelleted cells were solubilized in lysis buffer (2% SDS, 200 mM TEAB, 50 mM CAA, 10 mM TCEP) and boiled 5 minutes at 95°C. Protein concentration of the supernatant was estimated by BCA dosage. Thirty micrograms of each sample were digested using trypsin (Promega) and S-Trap Micro Spin Column was used according to the manufacturer’s protocol (Protifi, Farmingdale, NY, USA). Peptides were then speed-vacuum dried, then solubilized in 30 µl of 10 % acetonitrile (ACN) containing 0.1% TFA.

Liquid Chromatography-coupled Mass spectrometry analysis (nLC-MS/MS) analyses were performed on a Dionex U3000 HPLC nanoflow chromatographic system (Thermo Fischer Scientific, Les Ulis, France) coupled to a TIMS-TOF Pro mass spectrometer (Bruker Daltonik GmbH, Bremen, Germany). 2 µl were directly loaded on an Aurora C18 reverse phase resin (1.6 μm particle size, 100Å pore size, 75μm inner diameter, 25cm length mounted to the Captive nanoSpray Ionisation module, from IonOpticks, Middle Camberwell Australia) with a 4h run time with a gradient ranging from 98% of solvent A containing 0.1% formic acid in milliQ-grade H2O to 35% of solvent B containing 80% ACN, 0.085% formic acid in mQH2O.

The mass spectrometer acquired data throughout the elution process and operated in DIA PASEF mode with a 1.38 second/cycle, with Timed Ion Mobility Spectrometry (TIMS) enabled and a data-independent scheme with full MS scans in Parallel Accumulation and Serial Fragmentation (PASEF). Ion accumulation and ramp time in the dual TIMS analyzer were set to 100 ms each and the ion mobility range was set from 1/K0 = 0.63 Vs cm-2 to 1.43 Vs cm-2. Precursor ions for MS/MS analysis were isolated in positive polarity with PASEF in the 100-1.700 m/z range by synchronizing quadrupole switching events with the precursor elution profile from the TIMS device.

The mass spectrometry data were analyzed using DIA-NN version 1.8.1 (*79*). The database used for in silico generation of spectral library was a concatenation of Human sequences from the Swissprot database (release 2022-05) and a list of contaminant sequences from Maxquant and from the cRAP (common Repository of Adventitious Proteins). M-Terminus exclusion and carbamidomethylation of cysteins was set as permanent modification and one trypsin misscleavage was allowed. Precursor false discovery rate (FDR) was kept below 1%. The “match between runs” (MBR) option was allowed.

The results files of DIA-NN were analysed with Perseus version 1.6.15.0 (*80*). Contaminants and reverse proteins were eliminated. The data were transformed to Log 2 and only proteins with at least 2 valid values in each group and 70% of valid values in at least one group were kept. An unpaired two-sample test was applied to determine proteins with a q-value <0.05.

The raw data and extracted proteomics data have been deposited to the ProteomeXchange Consortium via the PRIDE (*81*) partner repository with the dataset identifier PXD066072.

### Proteomics – CK1α immunoprecipitation analysis

Control and CK1α sgRNA transduced THP1 cells (30×10^6^ cells) were then lysed in RIPA buffer (25 mM Tris HCl pH 7.4, 150 mM NaCl, 0.1% SDS, 0.5% Na-Deoxycholate, 1% NP-40, 1 mM EDTA) supplemented with 1X Halt Protease Inhibitor cocktail (ThermoFisher Scientific) and Benzonase (Novagen) as recommended by the manufacturer. Insoluble material was removed by centrifugation at 10,000*g*. For each Immunoprecipitation condition, 2.5 mg of lysate was precleared with Protein G Sepharose (Sigma) for 30 min, prior to incubation with 1 µg of CK1a antibody (A301-991A, Bethyl) and Protein G Sepharose for 2h at RT or overnight at 4°C under rotation. Samples were washed in lysis buffer four times before the addition of 2X Laemmli buffer. The Co-IP samples were solubilized volume by volume in lysis buffer containing 4% SDS, 400mM TEAB, pH 8.5, 20 mM TCEP, 100 mM chloroacetamide. Bottom-up experiments’ tryptic peptides were obtained by S-Trap Micro Spin Column according to the manufacturer’s protocol (Protifi, NY, USA). Briefly: Proteins were digested during 14h at 37°C with 1µg Trypsin sequencing grade (Promega). The S-Trap Micro Spin Column was used as described above, but eluted peptides were solubilized in 10 µl of 10% acetonitrile (ACN), 0.1% trifluoroacetic acid (TFA).

nLC-MS/MS analyses were performed with the same setup as described for the whole proteome above. One μL of peptides was loaded, concentrated, and washed for 3min on a C18 reverse phase Pepmap neo precolumn (3μm particle size, 300 μm inner diameter, 5 mm length, from Thermo Fisher Scientific). Peptides were then separated at 50°C on an Aurora C18 reverse phase resin (1.6 μm particle size, 100Å pore size, 75μm inner diameter, 25cm length) (IonOpticks, Middle Camberwell Australia) with a 60 minutes overall run-time gradient from 99% of solvent A containing 0.1% formic acid in milliQ-grade H2O to 40% of solvent B containing 80% ACN, 0.085% formic acid in mQH2O with a flow rate of 400 nL/min. The mass spectrometer acquired data throughout the elution process in a positive mode and operated in DIA PASEF mode with a 1.38 second/cycle, with Timed Ion Mobility Spectrometry (TIMS) enabled. Ion accumulation and ramp time in the dual TIMS analyzer were set to 100 ms each. The MS1 spectra were collected in the m/z range of 100–1,700. The diaPASEF window scheme ranged in dimensions from m/z 400 to 1,200 and in dimension 1/K0 from 0.65–1.42. The collision energy was set by linear interpolation between 59 eV at an inverse reduced mobility (1/K0) of 1.60 versus/cm2 and 20 eV at 0.6 versus/cm2.

The mass spectrometry data were analyzed as described above, except that normalization wasn’t allowed. The results files of DIA-NN were analyzed using a custom-made R-script (available at https://github.com/MarjorieLeduc/Shiny_PROTEOM_IC). For the comparison WT3/KO, a Welch T-test greater Unpaired were done on proteins showing at least 3 valid values in one group and at least 70% of valid values in the other group using log2(LFQ intensity).

Significant threshold is p-value<0.05. The raw data and extracted proteomics data have been deposited to the ProteomeXchange Consortium via the PRIDE (*81*) partner repository with the dataset identifier PXD066276.

### Proteomics - APEX2 proximitome analysis

The sequence coding for CSNK1A1 (obtained from addgene plasmid #123319) was tagged in N-terminus using APEX2 sequence (from addgene (Plasmid #49386), and cloned in the backbone plasmid pLVX-EF1a-IRES-Puro from ClonTech. THP-1 stable cell line expressing the pLVX-EF1a-IRES-Puro-APEX2-CSNK1A1 construct was established in the THP1 KO CSNK1A1 background (THP1-CK1α^sg2^ lacking endogenous CK1α) by lentiviral infection and puromycin selection (1μg/ml). Transgene expression and APEX2 activity (after biotin-phenol labeling) were verified by Western blot. Biotin-phenol labeling was performed in live cells after stimulation by 1μg/ml G3-YSD (DNA) for 2 hours or not, then biotinylated proteins from corresponding cell lysates were isolated using Streptavidin-coated magnetic beads (Pierce) as described in (*82*). Proteins captured on streptavidin-coated beads were digested on beads in 50μL of denaturing, reducing and alkylating buffer (0.1M TEAB, 10mM TCEP, 40mM Chloro-acetamide). Proteolysis took place overnight at 37°C under 1200 rpm shaking on thermomix with 1μg Trypsin sequencing grade (Sciex). After supernatant retrieval, peptides were desalted using Oasis HLB solid phase extraction device (hydrophilic N-vinylpyrrolidone and lipophilic divinylbenzene mixed polymer from Waters), then eluted, speed-vacuum dried and solubilized in 0.1% formic acid (FA).

nLC-MS/MS analyses were performed on a nanoElute 2 nanoflow system coupled to a TIMS-TOF Pro 2 mass spectrometer (Bruker Daltonik GmbH, Bremen, Germany). Two μL of the solubilized peptides were separated on an Aurora C_18_ reverse phase resin (1.7 μm particle size, 100Å pore size, 75 μm inner diameter, 25 cm length (IonOpticks, Middle Camberwell Australia) mounted onto the Captive nanoSpray Ionisation module, with a 60 minutes overall run-time gradient ranging from 99% of solvent A containing 0.1% formic acid in milliQ-grade H_2_O to 40% of solvent B containing 80% acetonitrile, 0.085% formic acid in mQH_2_O. nLC Eluate flow was electrosprayed through the captive spray module into the mass spectrometer (Bruker), which operated throughout the elution process in Data-Independent Analysis (DIA) mode, with PASEF-enabled method using the TIMS-Control software 5.1.8 (Bruker). The m/z acquisition range was 300–1201 with an ion mobility range of 0.75–1.25 1/K0 [V s/cm2], which corresponded to an estimated cycle time of 1.06 seconds at 100% duty cycle. DIA-PASEF windows, and collision energy were also left to default with a base of 0.85 1/K0 [V s/cm2] set at 20 eV and 1.3 1/K0 [V s/cm2] set at 59 eV. TOF mass calibration and TIMS ion mobility 1/K0 were performed linearly using three reference ions at 622, 922, and 1222 m/z (Agilent Technology, Santa Clara, CA, USA).

pLC-Flag-CSNK1A1 WT-Puro was a gift from Eva Gottwein (Addgene plasmid # 123319; http://n2t.net/addgene:123319;RRID:Addgene_123319).

pcDNA3 APEX2-NES was a gift from Alice Ting (Addgene plasmid # 49386; http://n2t.net/addgene:49386;RRID:Addgene_49386)

Identification and quantification data were extracted using Spectronaut® v.19.0.240606.62635 (Biognosys AG, Schlieren, Switzerland) as described in the setup.txt file available in the repository. Biefly: Spectronaut® DirectDIA+ (library-free “deep”) mode allowed generation of modelized peptides out of the human uniprot knowledge base organism_9606 (version 2024, april 3rd with 42 514 entries) in fasta format, along with the common contaminant database. The enzyme’s cleavage specificity was that of trypsin’s. The precursors and fragments’ mass tolerances were set to a maximum of +/-15 ppm variations. Modification of methionines by oxidation and acetylation of protein N-termini were set as possible events while all thiol groups from cysteines were considered completely thio-methylated. The software’s deep learning algorithm permitted an overall range of identification of over 4000 protein groups. Data were normalized between samples, match between runs was allowed to interpolate missing identifications. Data imputation was performed for ratio calculations for proteins represented in a minimum of 70% of samples. A threshold of adjusted p-value <0.01 was applied so the output (precursors) was applied. Retention time alignment and correction for mass accuracy were performed automatically. Spectronaut® statistical tools were used to visualize data, assess performances and quality before quantitatively compare groups of replicates’ datasets. The resulting proteins LFQ values were log2(x) transformed to reduce scale of protein intensities in the differential abundance analysis. We defined thresholds for up- and down-regulated proteins as absolute (log2Ratio) > 0.58 and adjusted p-value (or q-value) < 0.05. These ratios were defined to determine the level of modification of the protein expression. The raw data generated by DIA-PASEF and extracted proteomics data have been deposited to the ProteomeXchange Consortium via the PRIDE (*81*) partner repository with the dataset identifier PXD063855.

## Acknowledgments

We thank all members of the SOAP laboratory for their help and insightful discussions especially Célia Jaunasse; Johanna Bruce, Virginie Salnot Marjorie Leduc, and Emilie-Fleur Gautier (Proteom’IC facility, University Paris Cité, CNRS, INSERM, Institut Cochin, F-75014 PARIS, France) for the expertise and technical support for proteomic analysis; Aurélie Fétiveau and Séverine Marionneau-Lambot (CRCI^2^NA, Nantes, France) for technical assistance; the Cytocell and MicroPicell facilities (SFR Santé François Bonamy, Nantes, France) for the expertise; Floris Foijer (University of Groningen, Netherlands), Christina Woo (Harvard University, US), Daniel Krappmann (LMU, Germany), Joëlle Gaschet (CRCI^2^NA, Nantes, France) for providing essential tools. This work was supported by the DIM Thérapie Génique Paris Ile-de-France Région, IBiSA, and the Labex GR-EX.

## Funding

Ligue Nationale contre le Cancer, Equipe labellisée (JG, PJJ)

Ligue Nationale contre le Cancer, comités de Loire-Atlantique, Maine et Loire, Vendée, Ille et Vilaine, Mayenne, Finistère (JG, NB)

Institut National Du Cancer INCa_18384 (NB)

Institut National Du Cancer INCa PAIR-CEREB lNCa_16285 (JG)

Fondation ARC contre le Cancer PJA2021060003886 (NB) Fondation de France (JJ)

SIRIC ILIAD INCA-DGOS-INSERM-ITMO Cancer_18011 (JG)

## Author contributions

Conceptualization: JJ, KT, FG, SBN, NL, PPJ, JG, NB, GAG

Methodology: JJ, MT, KT, VJ, GAG, FG, SBN, LM, RM

Investigation: JJ, MT, KT, VJ, GAG, FG, LM, RM

Visualization: JJ

Supervision: SBN, NL, PPJ, NB

Writing—original draft: JJ, NB

Writing—review & editing: JJ, SBN, NL, PPJ, JG, NB

