## Supplemental Figures for "The Kinase CK1α coordinates Initiation and Termination of the cGAS-STING Pathway"

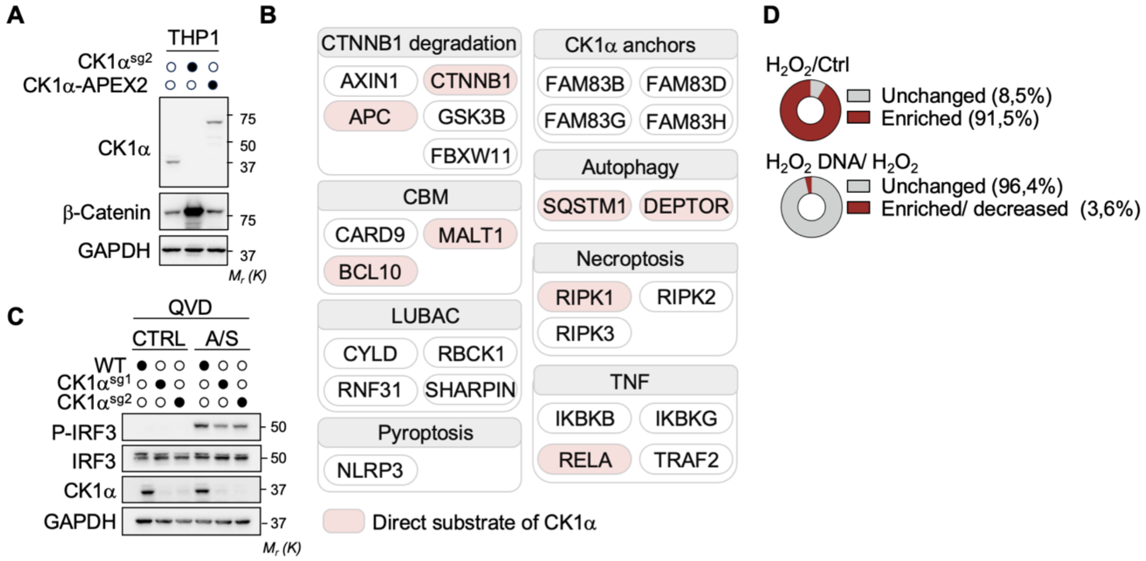


**Fig. S1. CK1α Proximity-dependent Labelling**

(**A**) THP-1 cells transduced with CK1α sgRNA were engineered to express the proximity-dependent labeling constructs CK1α-BirA and CK1α-APEX2. Cell lysates were prepared and analyzed by immunoblotting, as indicated. (**B**) Analysis of known physical protein interactors and substrates of CK1α that were significantly biotinylated in the CK1α-APEX2-based proximity proteome. Known CK1α interactors were sorted into the following categories: the CTNNB1 degradation complex, the Antigen receptor associated CARD protein-BCL10-MALT1 (CBM) complexes, the Linear ubiquitin chain assembly complex (LUBAC), the FAM83 family of CK1α anchors, autophagy, necroptosis, pyroptosis, and Tumor Necrosis Factor (TNF) signaling. (**C**) Immunoblots from THP-1 cells transduced with sgRNA against CK1α and co-treated with 10 µM of the BH3-mimetic compounds ABT-199 and S638445 for 4h. To prevent apoptosis, cells were pre-treated with 10 µM of the pancaspase inhibitor Q-VD-OPh for 15 Min prior to treatment with the BH3-mimetics. (**D**) Pie chart representing the percentage of CK1α proximity interactors enriched (or decreased) at steady state ($H_2O_2$/ Ctrl) or upon DNA stimulation ($H_2O_2$ DNA/ $H_2O_2$).

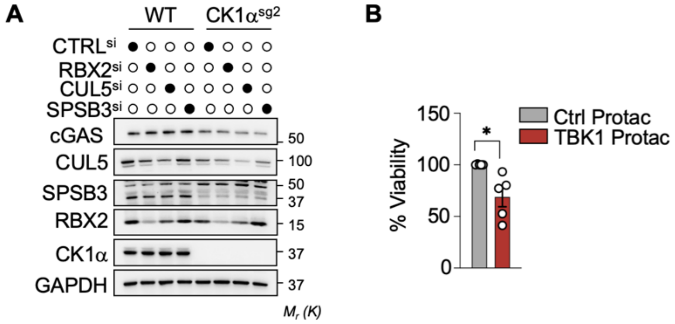


**Fig. S2. Identification of the CLR Complex Responsible for the steady-state Turnover of cGAS**

(**A**) cGAS protein abundance assessed by immunoblotting in sgRNA transduced THP-1 cells transfected with the indicated siRNA. (**B**) Cell viability of BT-549 cells treated with TBK1 or control PROTAC for (1µM, 48h) assessed by Celltiter Glo. Unpaired t-test, *p < 0.05, ** p < 0.01, *** p < 0.001, **** p < 0.0001.

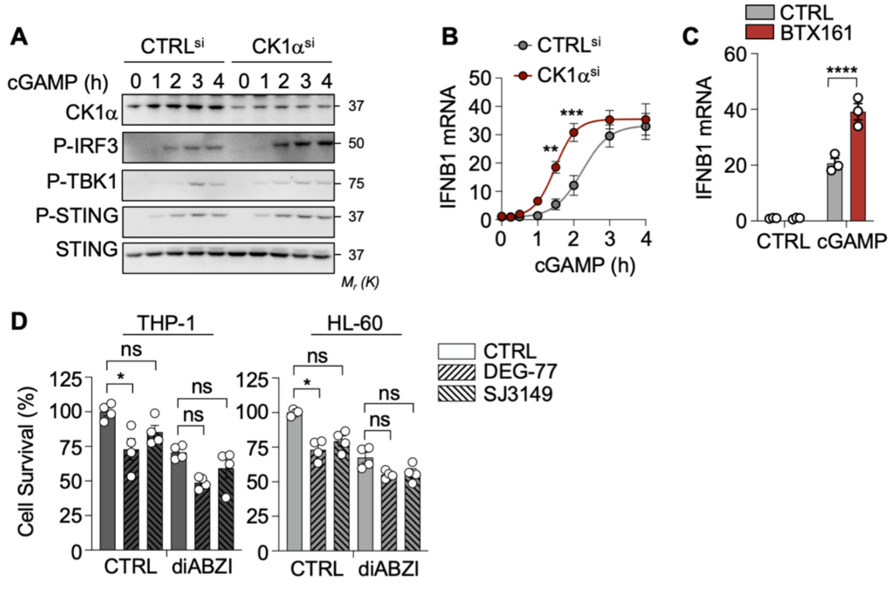


**Fig. S3. CK1α depletion potentiates STING signaling**

(**A**) Immunoblots of cell lysates and (**B**) IFNB1 expression quantified by RT-qPCR from THP-1 cells transfected with siRNA duplexes targeting CK1α and stimulated with the STING agonist 3'3'cGAMP (10 µg.ml$^{-1}$) for the indicated time. (**C**) IFNB1 expression quantified by RT-qPCR in THP-1 cells treated with the CK1α-targeted molecular glue BTX161 (10µM, 48h). (**D**) Cell viability assessed in p53-mutated AML cells (THP-1, HL-60) upon pre-treatment with CK1α degraders (100 nM DEG-77 or 50 nM SJ3149) for 4h and stimulation with diABZI at 5 nM (HL-60) or 200 nM (THP-1) for 16h. Ordinary two-way ANOVA, *$p < 0.05$, ** $p < 0.01$, *** $p < 0.001$, **** $p < 0.0001$.
